# Arsenic induces two different interaction modes of SUMO with promyelocytic leukemia (PML) proteins

**DOI:** 10.1101/2023.04.24.538205

**Authors:** Seishiro Hirano, Osamu Udagawa, Ayaka Kato-Udagawa

## Abstract

Promyelocytic leukemia-nuclear bodies (PML-NBs) are dot-like protein assemblies and implicated in the pathogenesis of leukemia and viral infection. PML is the scaffold protein of PML-NB and its client proteins such as SUMO, DAXX, and Sp100 reside in PML-NBs. It is known that a short exposure to trivalent arsenic (As^3+^) induces the solubility change and the subsequent SUMOylation of PML, and the SUMO interacting motif (SIM) is not necessary for these biochemical changes. However, it has not been well studied how As^3+^ initiates or enhances the association of SUMO with PML and the other PML-NB client proteins. Here, we report that As^3+^ enhanced non-covalent association of PML with SUMO via the SUMO-SIM interaction which is dispensable for the solubility change and SUMOylation of PML. We also report that the As^3+^-induced solubility change of PML was not affected by ML792, a SUMO E1 enzyme inhibitor, even though the nuclear localization of SUMO2/3 and protein SUMOylation were halted by ML792. As^3+^ did not change the solubility of DAXX and SUMOylation enzymes such as SAE1, UBA2, and UBC9. In contrast, As^3+^ induced SUMOylation of Sp100 with a concomitant loss of its solubility like PML in human leukemia cell lines. Our current results indicate that both covalent and non-covalent associations of SUMO with PML are increased in As^3+^-exposed cells, and Sp100 may play a role in the maintenance of PML-NBs.

## Introduction

Promyelocytic leukemia (PML) is a member of RBCC/tripartite motif (TRIM) family and a scaffold protein of PML-nuclear bodies (PML-NBs) [1, 2]. PML plays a role in the cell cycle, senescence, and tumor suppression [2], and is implicated in the pathogenesis of acute promyelocytic leukemia (APL) [3, 4]. However, the role of PML and PML-NBs *in vivo* are enigmatic because *Pml*^-/-^ mice grow normally [5], except that they are more vulnerable to viral infection [6, 7] and have less IFN-mediated angiostatic activity [8] than wild-type mice. The integrity of PML-NB is known to be modified by viral infection [9–11] and its components such as PML, Sp100, DAXX, and ATRX are important for signaling of both Type I and II interferons [12, 13]. PML may control the small ubiquitin-like modifier (SUMO) pool which is a critical regulator for human adenovirus replication [14].

Sp100 was first identified as a nuclear antigen recognized by autoantibodies obtained from primary biliary cirrhosis, and is found as nuclear dots (PML-NBs) in hepatic cells [15, 16]. Sp100 is an acidic protein harboring the NLS and shows a highly aberrant electrophoretic mobility in SDS-polyacrylamide gel electrophoresis [17]. Although the functional role of Sp100 is obscure, Sp100 proteins can homo-oligomerize and produce small-sized Sp100-containing foci in the nucleoplasm in the absence of PML [18]. Sp100 is present in both cytoplasmic and nuclear regions. Sp100 is modified with SUMO1 upon IFN-treatment and the modified Sp100 is extractable as a nuclear component [19]. The DAXX-ATRX complex plays a key role in the recruitment of histone H3.3, and PML-NBs regulate the DAXX-ATRX-H3.3 axis [20, 21]. It has been reported that the loss of PML shifts the histone H3 methylation balance from H3K9me3 to H3K27me3 [20, 22, 23]. Recently, it has been reported that DAXX functions as a disaggregase and an unfoldase. DAXX disassembles ATXN1(82Q) aggregates and Aβ42 fibrils into a soluble state, and solubilizes p53 and MDM2 molecules from their aggregated forms [24].

DAXX has SUMO-interacting motifs (SIMs) on both N- and C-terminals which are linked with the intrinsically disordered region (IDR) [25]. Both DAXX and Sp100 co-localize with PML in PML-NBs through an interaction of SIMs of these proteins with SUMOylated PML at K160. However, the SUMO-SIM interaction is not required for PML to maintain the lattice formation of PML-NBs [26]. The SIM at the C-terminal of DAXX is required for localization of DAXX and ATRX in PML-NBs [27].

Arsenite (As^3+^) is known to degrade oncogenic PML-retinoic acid receptor α fusion proteins (PML-RARα) via the SUMOylation-dependent pathway and restore normal PML-NBs, replacing PML-RARα small speckles in APL cells [28–31]. The solubility loss and SUMOylation of PML are two overt biochemical changes in As^3+^-treated cells [32–34]. However, it remained to be elucidated why PML is insolubilized and SUMOylated upon exposure to As^3+^. To our best knowledge, the specific and rapid biochemical changes of PML are caused by As^3+^ and antimony (Sb^3+^), but not by the other toxic metals [32, 35]. Although the biochemical modification of PML is reported to be caused by oxidative stress [36], As^3+^ induces the solubility change of PML without an overt activation of antioxidant responsive element (ARE) [34].

It is well documented that the As^3+^-induced solubility change and SUMOylation of PML are completed in a short time [32-34, 37, 38]. However, little is known about how As^3+^ initiates the association of SUMO with PML, and whether the other PML-NB partner proteins are changed by As^3+^. We report that As^3+^ increases a non-covalent association of PML with SUMO molecules via the SUMO-SIM interaction which is not requisite for the solubility loss and the covalent SUMOylation of PML. Sp100, but not the other PML-NB client proteins and SUMOylation enzymes responds to As^3+^ in the similar fashion as PML.

## Materials and Methods

### Chemicals, antibodies, plasmids, and primers

Sodium *m*-arsenite and IFNα2a were purchased from Sigma-Aldrich (St. Louis, MO). ML792 (SUMO E1 inhibitor) and TAK243 (ubiquitin E1 inhibitor) were purchased from MedKoo Bioscience (Morrisville, NC) and Selleckchem (Houston, TX), respectively. A BCA protein assay kit was obtained from Pierce-ThermoFisher (Rockford, MA). Alexa Fluor™ 488-tagged anti-SUMO2/3 antibody was labelled using an Alexa Fluor^®^ 488 Antibody Labeling Kit (Invitrogen-ThermoFisher, Waltham, MA). Unless otherwise specified, common chemicals of analytical grade were obtained from Sigma-Aldrich or Fujifilm-Wako (Osaka, Japan). Details of antibodies, custom-made plasmids, and PCR primers are provided in the Supporting information.

### Cells

HEK293 and HL60 cells were obtained from RIKEN (Ibaraki, Japan), and Jurkat cells (ECACC authorized) were obtained from DS Pharma Biomedical (Osaka, Japan). PML double nickase plasmid (Santa Cruz, Dallas, TX) was transduced into HEK293 cells using Lipofectamine LTX (Invitrogen-ThermoFisher) to knock out *PML* by the CRISPR-Cas9 system. The stable transfectants were selected using puromycin and designated as *PML^-/-^*cells. HEK293, which were stably transfected with the human *PML-VI* gene (RC220236, OriGene, Rockville, MD), were selected using neomycin and were designated as HEKPML cells [34]. HEK293 and HEKPML cells were further transduced with mCherry-tagged SUMO2 plasmid vectors (Wild, K11R, and C-terminal ΔGG) and stable transfectants were selected using puromycin. The live cell images were captured by inverted fluorescent microscopy (TS100, Nikon, Tokyo).

### Western blotting

Cells were lysed with ice-cold RIPA buffer (Santa Cruz, Dallas, TX) containing 0.1% sodium dodecyl sulfate, protease inhibitors (Santa Cruz) and phosphatase inhibitor cocktails (Calbiochem/Merk Millipore, San Diego, CA) on ice for 10 min. The lysate was centrifuged at 9,000 g for 5 min at 4 °C. The supernatant was labelled as the RIPA-soluble fraction (Sol). The pellet was rinsed with PBS and suspended in150 mM Tris-HCl buffer (pH 7.2) of the same volume as the RIPA buffer containing Benzonase^®^ nuclease (250 U/mL, Santa Cruz). The sample was incubated at 25 °C for 2 h with intermittent mixing (Thermomixer comfort, Eppendorf, Wesseling-Berzdorf, Germany) to reduce viscosity. The digested sample was labelled as the RIPA-insoluble fraction (Ins). Proteins were resolved by LDS (SDS)-PAGE and electroblotted onto PVDF membranes. Unless otherwise specified, NuPAGE 4-12% Bis-Tris gels (Invitrogen-ThermoFisher) were used for the electrophoresis. The membrane was blocked with PVDF Blocking Reagent (TOYOBO, Osaka, Japan) before probing with antibodies. The membrane was soaked in ECL (Prime, GE Healthcare, Buckinghamshire, UK) and signals were captured with CCD cameras (Lumino Imaging Analyzer, FAS-1100, TOYOBO; Amersham ImageQuant 800, GE Healthcare, Uppsala, Sweden). Antibodies were diluted with Can-Get-Signal solution (TOYOBO). The membrane was finally re-probed with HRP-conjugated anti-tubulin and HRP-conjugated anti-histone H3 antibodies to evaluate loading of cytosolic and nuclear proteins and distribution of these proteins into the soluble and insoluble fractions.

### Immunoprecipitation

The clear RIPA-soluble fraction was used for immunoprecipitation study. The protein concentration of the supernatant was adjusted to 2 mg/mL with RIPA. The sample was diluted with the same volume of 150 mM Tris-HCl solution (pH 7.2) and reacted with anti-SUMO1 or anti-SUMO2/3 affinity beads, or anti-RFP mAb-magnetic beads at 4-8 °C for 2 h with intermittent vortexing. The beads were washed three times with 150 mM Tris-HCl solution (pH 7.2) containing 0.1% NP40, and the pellet was extracted with 2X reducing agent-free LDS (SDS)-PAGE sample buffer at 30℃ for 10 min. The extracts were heated in the presence of 50 mM DTT at 95°C for 5 min to prepare the final samples for electrophoresis and the following western blotting.

### RT-qPCR

*DAXX* mRNA levels of HEK293, *PML^-/-^*, and HEKPML cells were measured by RT-qPCR adopting the ΔΔCt method. Briefly, total RNA was extracted from the cells using Trizol^®^ (ThermoFisher) and PureLink^®^ RNA Mini kit (ThermoFisher) with on-column DNase-treatment. Each cDNA was synthesized using a PrimeScript RT reagent kit (Takara Bio, Kusatsu-Shiga, Japan). Quantitative PCR reactions were performed using SYBR Premix Ex Taq II with a thermal cycler (TP800, Takara Bio).

### Immunofluorescent staining

HEK cells were grown in an 8-well chamber slide (Millicell EZ slide, Merck-Millipore, Burlington, MA) to early confluence. The chamber slide was pre-treated with 100 μg/mL Type VI collagen overnight in a refrigerator to enhance adhesion and spreading of the cells. Jurkat cells were cytocentrifuged at 1200 rpm for 3 min. The cells on the glass-slide were rinsed in warmed (37 °C) HBSS, fixed with 3.7% formaldehyde solution for 10 min, permeabilized with 0.1% Triton X-100 for 10 min, and treated with Image-iT^®^ FX Signal Enhancer (ThermoFisher) for another 10 min. Then, the cells were immunostained with Alexa Fluor^®^ 488-labeled anti-SUMO2/3 for 45 min. HEK293 cells were also immunostained with anti-PML polyclonal antibody followed by Alexa Fluor™ 594-labeled goat anti-rabbit IgG antibody before staining for SUMO2/3. Jurkat cells were stained with polyclonal anti-Sp100 antibody followed by Alexa Fluor™ 488-labeled goat anti-rabbit IgG antibody before staining with monoclonal anti-PML or monoclonal anti-SUMO2/3 antibody with Alexa Fluor™ 594-labeled goat anti-mouse IgG antibody. The antibodies were diluted in Can-Get-Signal^®^ immunostain solution A (TOYOBO) to 1/125, 1/200, and 1/1000 for primary antibodies, Alexa Fluor^®^ 488-labelled anti-SUMO2/3, and Alexa Fluor^®^-labeled secondary antibodies, respectively. The nuclei were counter-stained with DAPI. The fluorescence images were captured by fluorescence microscopy (Eclipse 80i, Nikon). The digital images were assembled using Adobe PHOTOSHOP^®^ and ImageJ (http://imagej.nih.gov/ij) software.

### Data analyses

The size of PML-NBs was evaluated by the average diameter of SUMO2/3 dots in the nuclei assuming the dots are spherical using ImageJ Software. The number and size of PML-NBs were presented as means ± SD. Densitometric data of western blotting and normalized *DAXX* mRNA levels were presented as means ± SEM. Statistical analyses were performed by one-way or two-way ANOVA followed by Tukey’s *post-hoc* test.

## Results

### Solubility changes and SUMOylation of PML following exposure to As^3+^ in HEK cells

It is known that human cells express six nuclear (I-VI) and one cytosolic (VII) PML isoforms [7]. Exposure to As^3+^ for 2 h changed the solubility and the electromobility of PML isoforms in HEK293 cells, and the mobility retardation was accompanied by SUMOylation of PML (Fig. 1A). The endogenous PML was detected as small PML-NBs in HEK293 cells and SUMO2/3 was co-localized with PML irrespective of exposure to As^3+^ (Fig. 1B), suggesting that the microscopically observable co-localization of PML and SUMO does not necessarily reflect the covalent SUMOylation of PML. Next, we analyzed the number and size of PML-NBs as SUMO2/3 dots quantitatively (Fig. 1C). The number of dots was significantly smaller, and the size was significantly larger in HEKPML than in HEK293 cells. Exposure to As^3+^ did not change the number of SUMO2/3 dots, and the size was only slightly but significantly (p value = 0.018) changed. Unlike HEK293 and HEKPML cells, SUMO2/3 only diffusely distributed over the nuclei in *PML* knockout (*PML^-/-^*) cells (Fig. 1C).

**Fig. 1.**
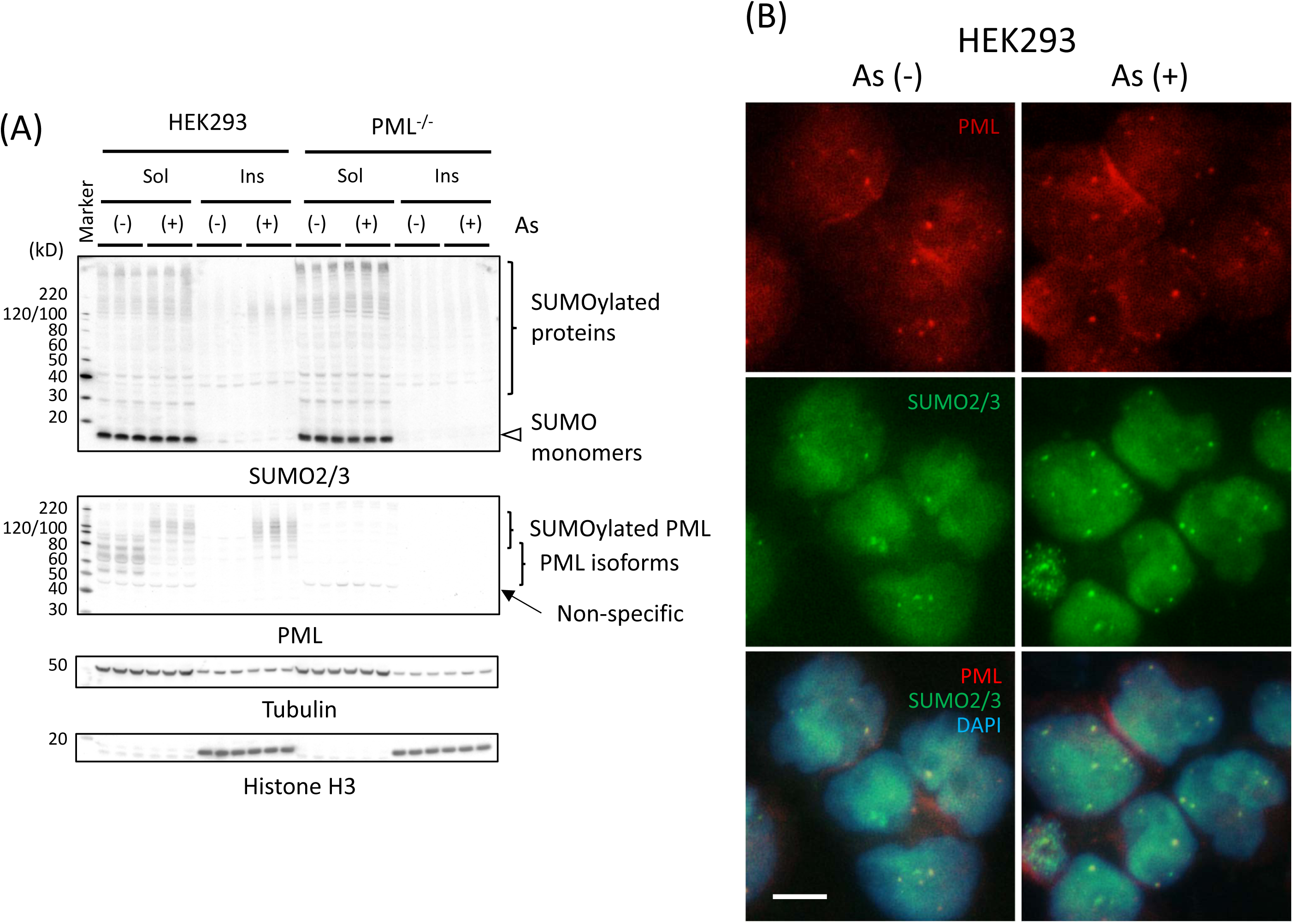
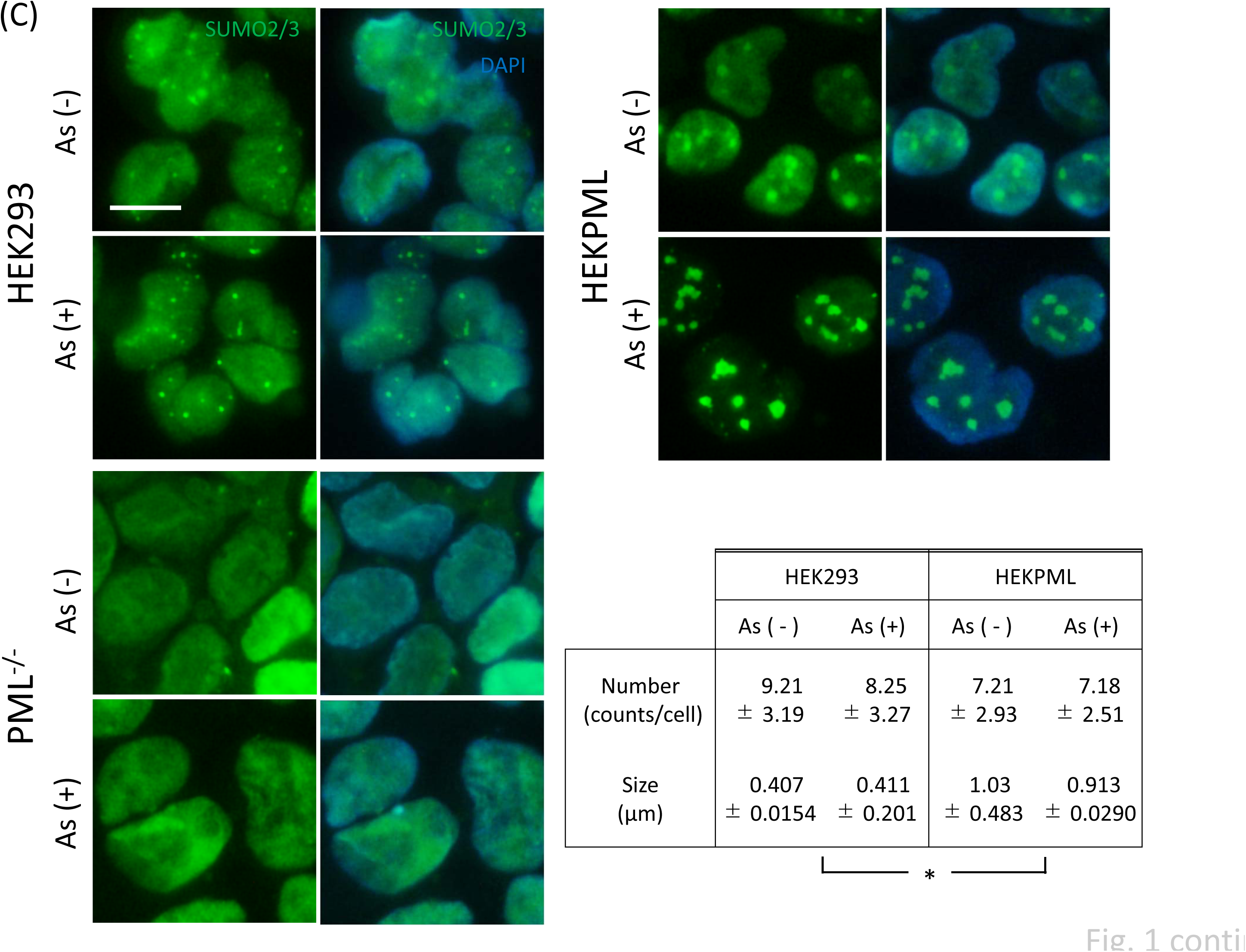
Immunoblot analyses of SUMO2/3 and PML in HEK293 and *PML^-/-^* cells (A), immunostaining of HEK293 cells for the detection of PML and SUMO2/3, and characterization of SOMO2/3 dots in HEK293, *PML^-/-^*, and HEKPML cells (C). (A) The cells were treated with 3 μM As^3+^ (+) or left untreated (-) for 2 h. The soluble (Sol) and insoluble (Ins) fractions of cell lysates were obtained from three different wells. The membrane was probed sequentially with anti-SUMO2/3, anti-PML, and finally with HRP-tagged anti-tubulin and anti-histone H3 antibodies. Note that the polyclonal anti-PML antibody detected endogenous PML isoforms in HEK293 cells. PMLs in the soluble fraction were partially shifted to the insoluble fraction, and SUMOylated by 2-h exposure to 3 μM As^3+^ in HEK293 cells. An open arrowhead indicates SUMO2/3 monomers. (B) The cells were treated with 3 μM As^3+^ for 2 h or left untreated before fixation and immunostaining with anti-PML and anti-SUMO2/3 antibodies. Note that small PML dots co-localized with SUMO2/3 dots in the nuclei of HEK293 cells. Scale bar = 10 μm. (C) HEK293, HEKPML, and *PML^-/-^* cells were treated with 3 μM As^3+^ or left untreated for 1 h. The cells were fixed, permeabilized, and stained with Alexa Fluoro 488-labeled anti-SUMO2/3 antibody. The nuclei were counterstained with DAPI. Note that SUMO2/3 was stained diffusely over the nuclei of *PML^-/-^* cells, and the nucleoplasmic PML (PML-VI) was more restricted to PML-NBs in As^3+^-exposed HEKPML cells. The size of PML-NBs (SUMO2/3 dots) was assayed by the estimated diameter assuming that the PML-NBs were spherical. Measurements were performed for more than 200 dots (201 – 212 dots) using ImageJ and presented as means ± SD. *, Both the number and the size of SUMO dots were significantly different between HEK293 and HEKPML cells. The size was slightly, but significantly different between untreated (-) and As^3+^-exposed (+) cells (p value = 0.018) as determined by two-way ANOVA.

### SUMO E1 enzyme is requisite for the nuclear localization of SUMO and PML SUMOylation

We used HEKPML cells, which overexpress PML-VI, for this experiment to observe the biochemical changes of PML and intracellular distribution of SUMO clearly. The SUMOylation with SUMO1 and SUMO2/3 was inhibited in the presence of ML792. In contrast, ubiquitinated proteins almost disappeared in TAK243-treated HEKPML cells regardless of 1 h-exposure to As^3+^ (Fig. 2A). It is of interest to note that SUMOylated proteins at the high molecular weight region (right half brackets) remained in the sample of TAK243-treated cells, suggesting that proteins migrating at these regions are degraded by SUMO-targeted ubiquitin E3 ligases (STUbL) and proteasomes [39] in the absence of the inhibitor. The localization of SUMO2/3 at PML-NBs was lost, and SUMO2/3 distributed diffusively in the cell including cytoplasm in ML792-treated HEKPML cells (Fig. 2B).

**Fig. 2.**
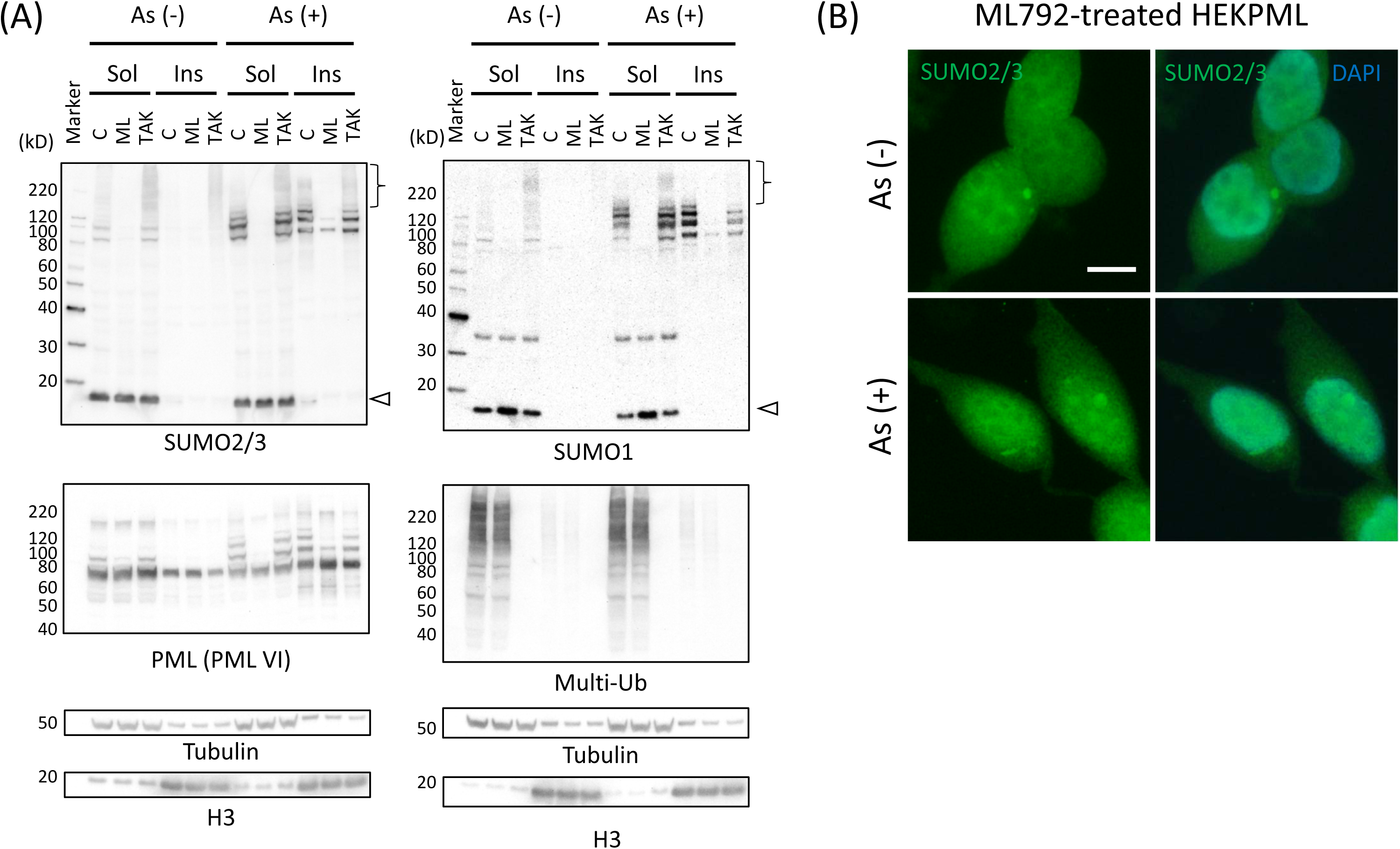
Effects of ML792 and TAK243 on SUMOylation and ubiquitination of proteins (A), and effects of ML792 on the cellular distribution of SUMO2/3 (B) in HEKPML cells. (A) HEKPML cells were pre-treated with 20 μM ML792 (SAE SUMO E1 inhibitor), 10 μM TAK243 (UAE ubiquitin E1 inhibitor) or 0.1% DMSO (vehicle, (C)) for 4 h, and further cultured in the presence (+) or absence (-) of 3 μM As^3+^ for 1 h. Immunoblot analyses were performed using anti-SUMO1, anti-SUMO2/3, anti-multi-ubiquitin, and anti-PML antibodies. An open arrowhead indicates SUMO1 or SUMO2/3 monomers. The right half parenthesis indicates SUMOylated proteins at high molecular weight regions. (B) Immunostaining of HEKPML cells with Alexa Fluoro 488-labeled anti-SUMO2/3 antibody. The cells were pre-treated with 20 μM ML792 for 4h and further cultured with and without 1-h exposure to 3 μM As^3+^. Scale bar = 10 μm.

### Expression of mCherry-tagged SUMO2 mutants in HEK cells and As^3+^-induced association of PML with SUMO2

Wild-type (Wild), K11R (polySUMOylation incompetent), and diglycine-deleted (ΔGG, SUMOylation incompetent) mCherry-conjugated SUMO2 were stably expressed in HEK293 and HEKPML cells. The solubility shift and SUMOylation of PML were observed similarly in the three different mCherry-conjugated SUMO2-expressing HEK293 cells after 2 h-exposure to 3 μM As^3+^ (Fig. 3). As we expected, conjugation of proteins with mCherry-SUMO2 was negligible in ΔGG mCherry-SUMO2-transduced HEK293 cells (a right half bracket).

**Fig. 3.**
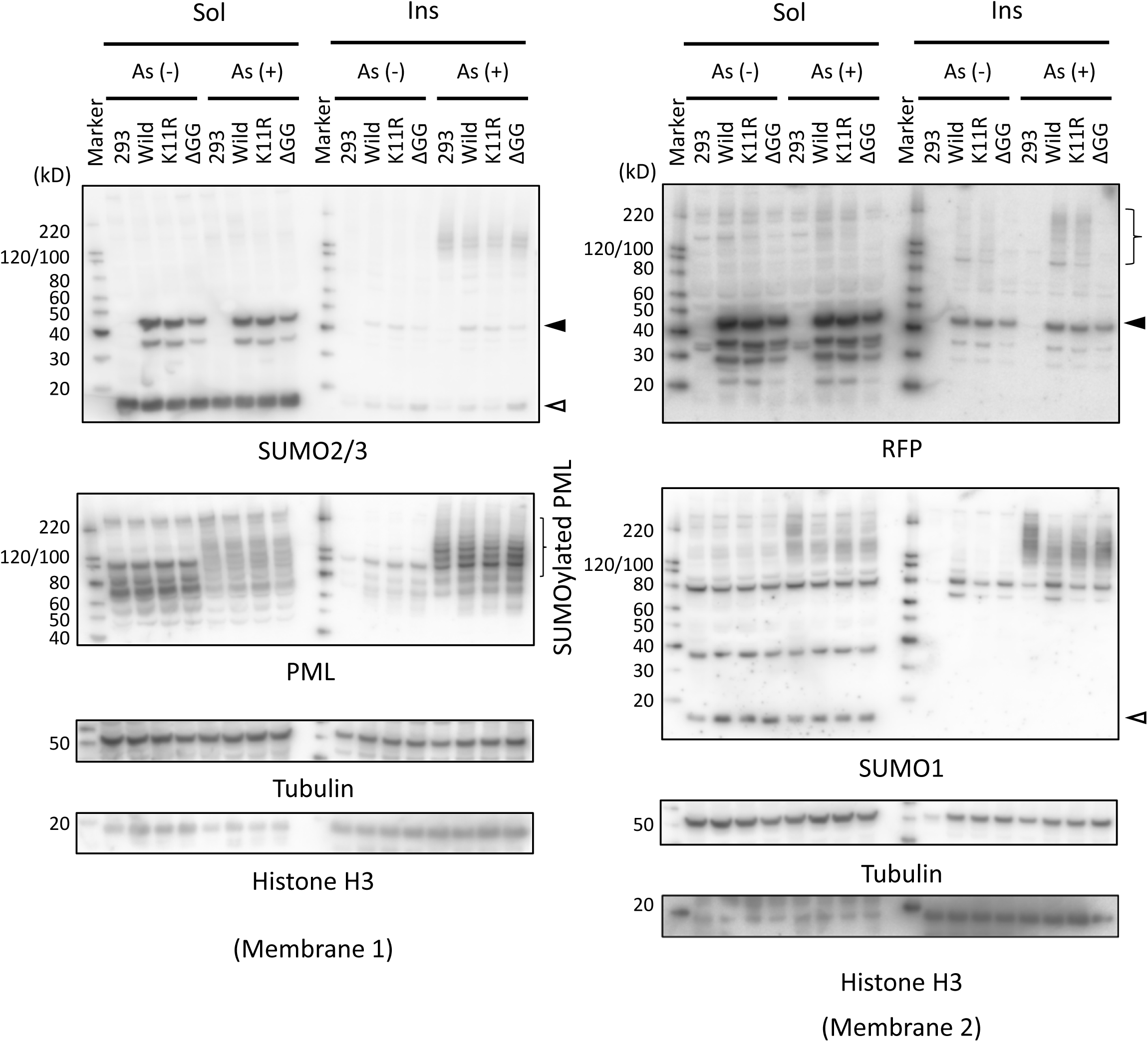
Immunoblot analyses of HEK293 cells stably expressing mCherry-conjugated wild-type and mutant SUMO2. (A) N-terminal mCherry expression plasmids harboring wild, K11R mutant (K11R), and C-terminal GG-deleted (ΔGG) *SUMO2* were stably transduced in HEK293 cells. The cells were treated with 3 μM As^3+^ for 2 h (As (+)) or left untreated (As (-)). The soluble (Sol) and insoluble (Ins) fractions of each cell lysate were analyzed for SUMO1, SUMO2/3, PML, and RFP by western blotting. Closed and open triangles indicate mCherry-conjugated SUMO2 monomer and endogenous SUMO monomer, respectively. 293, non-transfected HEK293 cells. The right half parenthesis indicates SUMOylated proteins with mCherry-conjugated SUMO2.

About a half amount of PML isoforms still remained in the soluble fraction after exposure to 3 μM As^3+^ for 1 h (Fig. 4A). Then, the clear soluble fractions were obtained from mCherry-SUMO2-transduced cells after 1 h-exposure to 3 μM As^3+^ for immunoprecipitation assay using anti-RFP antibody-conjugated magnetic beads. It is interesting that endogenous PML isoforms and SUMOylated PML were immunoprecipitated with both mCherry-tagged wild and ΔGG SUMO2 in As^3+^-exposed HEK293 cells (Fig. 4B). We further examined whether the SUMO-SIM interaction was required for the association of PML and SUMOylated PML with mCherry-SUMO2 in As^3+^-exposed cells using HEKPML cells. ΔGG mCherry-SUMO2 did not localize in PML-NBs in HEKPML cells (Fig. 5A). HEKPML cells express DDK(FLAG)-tagged PML-VI, and this PML isoform lacks a SIM motif. In contrast to HEK293 cells (Fig. 4B), PML-VI was not pulled down by anti-RFP antibody in mCherry-tagged ΔGG SUMO2 expressing HEKPML cells (Fig. 5B). These results indicate that non-covalent association of PML with SUMO2 was increased by As^3+^ via the SUMO-SIM interaction. The As^3+^-induced SUMOylation (covalent) and non-covalent interaction of PML with SUMO is schematically depicted in Fig. 6.

**Fig. 4.**
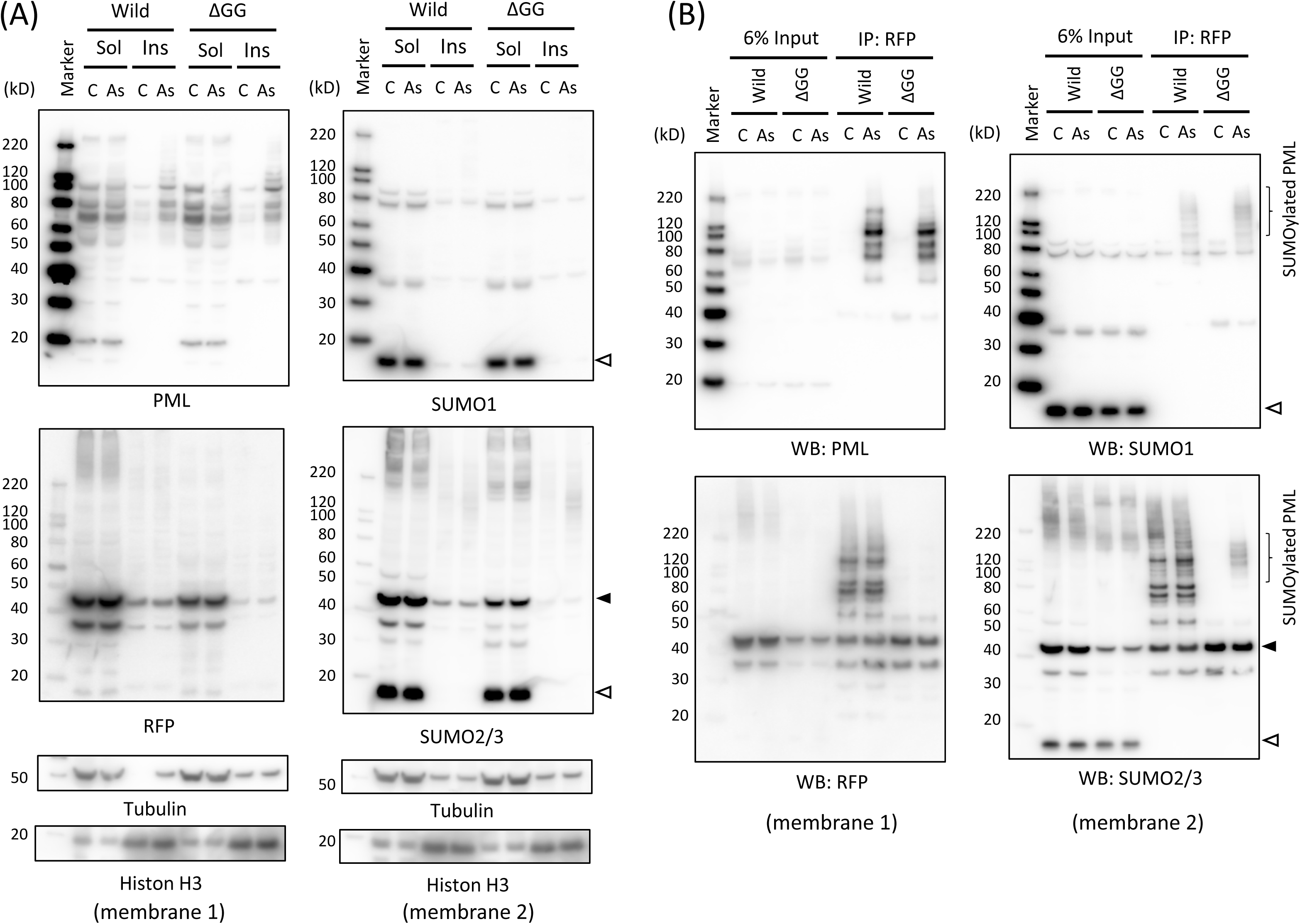
Immunoprecipitation analysis of HEK293 cell lysate with anti-RFP antibody. HEK293 cells stably expressing mCherry-conjugated SUMO2 (Wild) or mCherry-conjugated SUMO2 with C-terminal diglycine deletion (ΔGG) were used. The cells were treated with 3 μM As^3+^ (As) or left untreated (C) for 1 h and lyzed with RIPA buffer. (A) The cell lysates were separated into the soluble (Sol) and insoluble fractions (Ins) and analyzed for PML, SUMO1, SUMO2/3, and mCherry-conjugated SUMO2 (RFP) by western blotting. (B) The clear supernatants of the lysates were used for immunoprecipitation assay using anti-RFP-conjugated magnetic beads. Note that PML and SUMOylated PML were pulled down by ΔGG mCherry-SUMO2, suggesting that PML and PML SUMOylated with endogenous SUMO molecules associated with SUMO2 non-covalently in As^3+^-exposed HEK293 cells. An open triangle indicates endogenous SUMO1 or SUMO2/3 monomers. Closed and open triangles indicate mCherry-conjugated SUMO2 monomer and endogenous SUMO monomer, respectively.

**Fig. 5.**
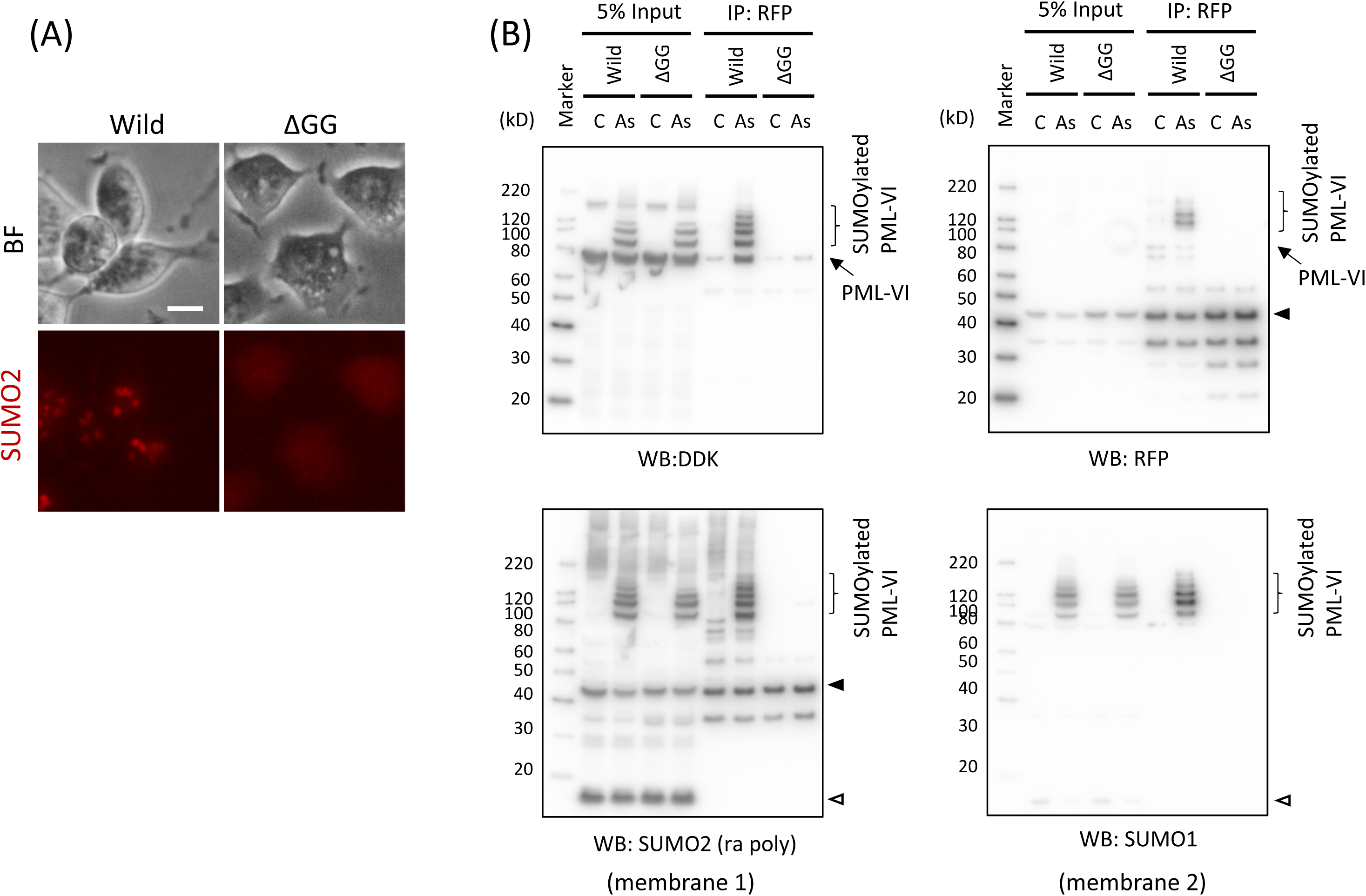
Microscopic observation of HEKPML cells expressing wild-type (Wild) or C-terminal diglycine-deleted (ΔGG) mCherry-SUMO2 (A), and immunoprecipitation analysis of HEKPML cell lysate with anti-RFP antibody (B). (A) The wild and ΔGG SUMO2 expressing HEKPML cells were cultured in a 48-well dish pre-treated with 100 μg/mL Type IV collagen. The live cell images of mCherry-SUMO2 and bright field (BF) were captured by fluorescence microscopy (Eclipse TS100, Nikon; Excitation filter 510-560 nm and Barrier filter 590 nm cut-on). Scale bar = 10 μm. (B) HEKPML cells stably expressing DDK(FLAG)-tagged PML-VI and mCherry-conjugated SUMO2 (Wild) or mCherry-conjugated SUMO2 with C-terminal diglycine deletion (ΔGG) were used. The cells were treated with 3 μM As^3+^ (As) or left untreated (C) for 1 h and lyzed with RIPA buffer. Note that the expression of DDK-tagged PML-VI was so high that endogenous PML isoforms were not visible. See also the legend to Fig. 4 for the immunoprecipitation procedure. PML-VI and SUMOylated PML-VI was not pulled-down by ΔGG mCherry-SUMO2, suggesting that non-covalent association did not occur between SUMO2 and PML-VI. Closed and open triangles indicate mCherry-conjugated SUMO2 monomer and endogenous SUMO monomer, respectively.

**Fig. 6.**
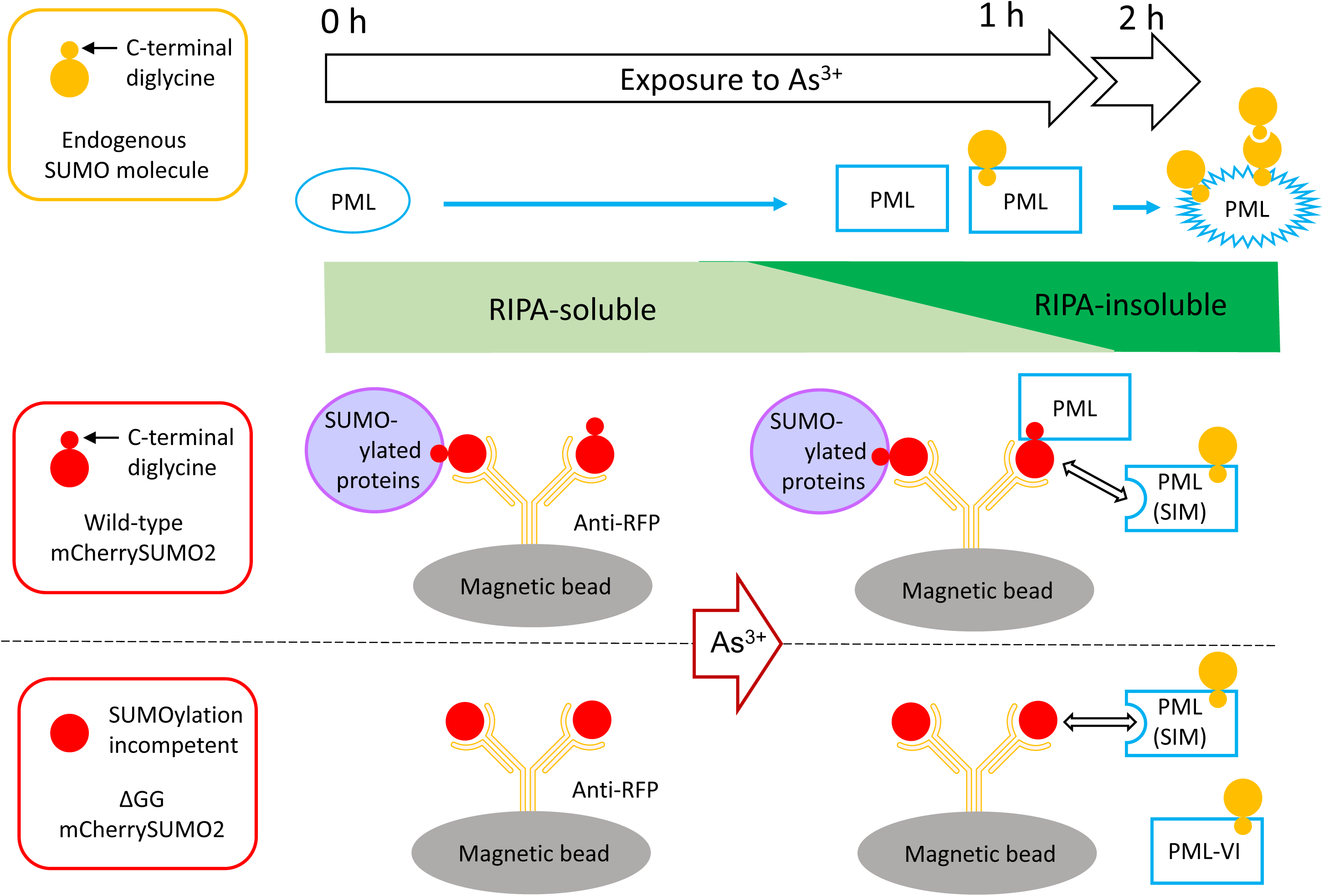
Schematic presentation for the pull-down assay using mCherry-conjugated SUMO2 (wild or ΔGG) expressing HEK293 cells. The solubility of PML in the RIPA buffer is lost gradually and PML is SUMOylated upon exposure to As^3+^ in HEK293 cells. This process appears to be completed in 2h. SUMOylated proteins including PML were pulled down by anti-RFP antibody-conjugated magnetic beads. In addition, PML and PML SUMOylated with endogenous SUMO were pulled down by via the SUMO-SIM (SUMO interacting motif) interaction in As^3+^-exposed cells. PML-VI, which does not have a SIM, is SUMOylated upon As^3+^-exposure like other PML isoforms. However, PML-VI cannot associate with SUMO2 molecules non-covalently.

### DAXX and SUMOylation enzymes in HEK cells

DAXX is one of the well-defined PML-NB client proteins [33, 40]. However, the response of DAXX to As^3+^ was different from that of PML, and the solubility of DAXX was not changed by As^3+^ (Fig. 7A). An ancillary but interesting finding is that the expression level of DAXX varied among the cell types. The protein level of DAXX in HEKPML cells was significantly lower than that in both HEK293 and *PML^-/-^* cells as examined using two different antibodies (Fig. 7B). However, the mRNA level of *DAXX* was not co-related to its protein level (Supplementary Fig. 1). Exposure to As^3+^ did not change the solubility of SUMOylation enzymes such as SAE1, UBA2, and Ubc9 (Fig. 7A).

**Fig. 7.**
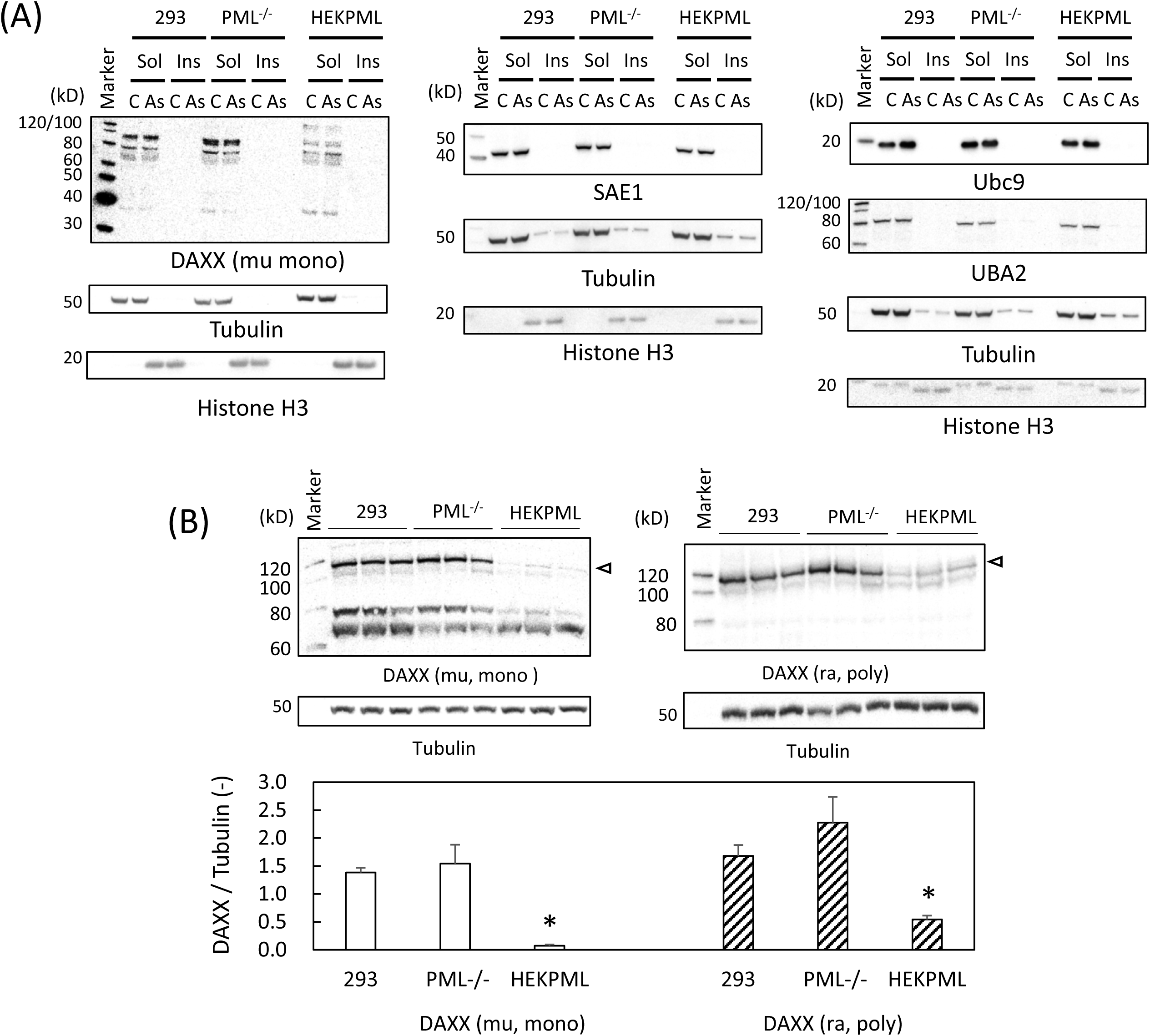
Immunoblot analyses of DAXX, SAE1, Ubc9, and UBA2 in HEK293 (293), *PML^-/-^*, and HEKPML cells. (A) Sub-confluent cells were exposed to 3 μM As^3+^ (As) or left untreated (C) for 2 h before lysis with cold RIPA buffer. The lysate was separated into the RIPA-soluble (Sol) and -insoluble (Ins) fractions for the detection of DAXX, SAE1, Ubc9, and UBA2 by immunoblotting. (B) The soluble fractions of cell lyzates were used for the quantitation of DAXX in untreated HEK293, *PML^-/-^*, and HEKPML cells in triplicate wells. The electrophoresis was performed using NuPAGE 3-8% Tris-Acetate gels (Invitrogen-ThermoFisher), and two different antibodies were used for the densitometric quantitation of the clearly separated DAXX band. An open triangle indicates protein bands used for the densitometric quantification. *, Significantly different from the other two groups (N=3).

### Sp100 and the other PML-NB proteins in Jurkat and HL60 cells

Unfortunately, Sp100 was not detectable in HEK cells by our current immunoblot method irrespective of the level of PML expression and exposure to As^3+^ even when the chemiluminescence signal was intensified (Supplementary Fig. 2A). The basal SUMOylation capacity with SUMO1 is not responsible for the undetectable level of Sp100 in HEK cells because RanGap1 is SUMOylated normally in HEK cells (Supplementary Fig. 2B) [17, 38]. Then, we used other human cell lines such as Jurkat and HL60 which express Sp100 moderately. As^3+^ changed Sp100 from the RIPA-soluble to RIPA-insoluble form and increased the molecular weight by about 12 kD in Jurkat cells (Fig. 8A). In contrast, the substantial amount of modified Sp100 was present in the soluble fraction of untreated HL60 cells (Fig. 8B). Sp100 showed the similar solubility change after exposure to As^3+^ in HL60 cells as in Jurkat cells (Fig. 8 and Supplementary Fig. 3A). The other PML-NB proteins of Jurkat and HL60 cells such as PML, SUMO, Ubc9, and DAXX responded to As^3+^ similarly to those of HEK cells (Fig. 8, and Supplementary Fig. 3). Sp100 co-localized with both SUMO2/3 and PML irrespective of As^3+^ exposure in the nuclei of Jurkat cells (Fig 8C). However, the immunoprecipitation study using anti-SUMO1, and anti-SUMO2/3 antibodies indicated that Sp100 was preferentially conjugated with SUMO1 in Jurkat and HL60 cells (Fig. 9).

**Fig. 8.**
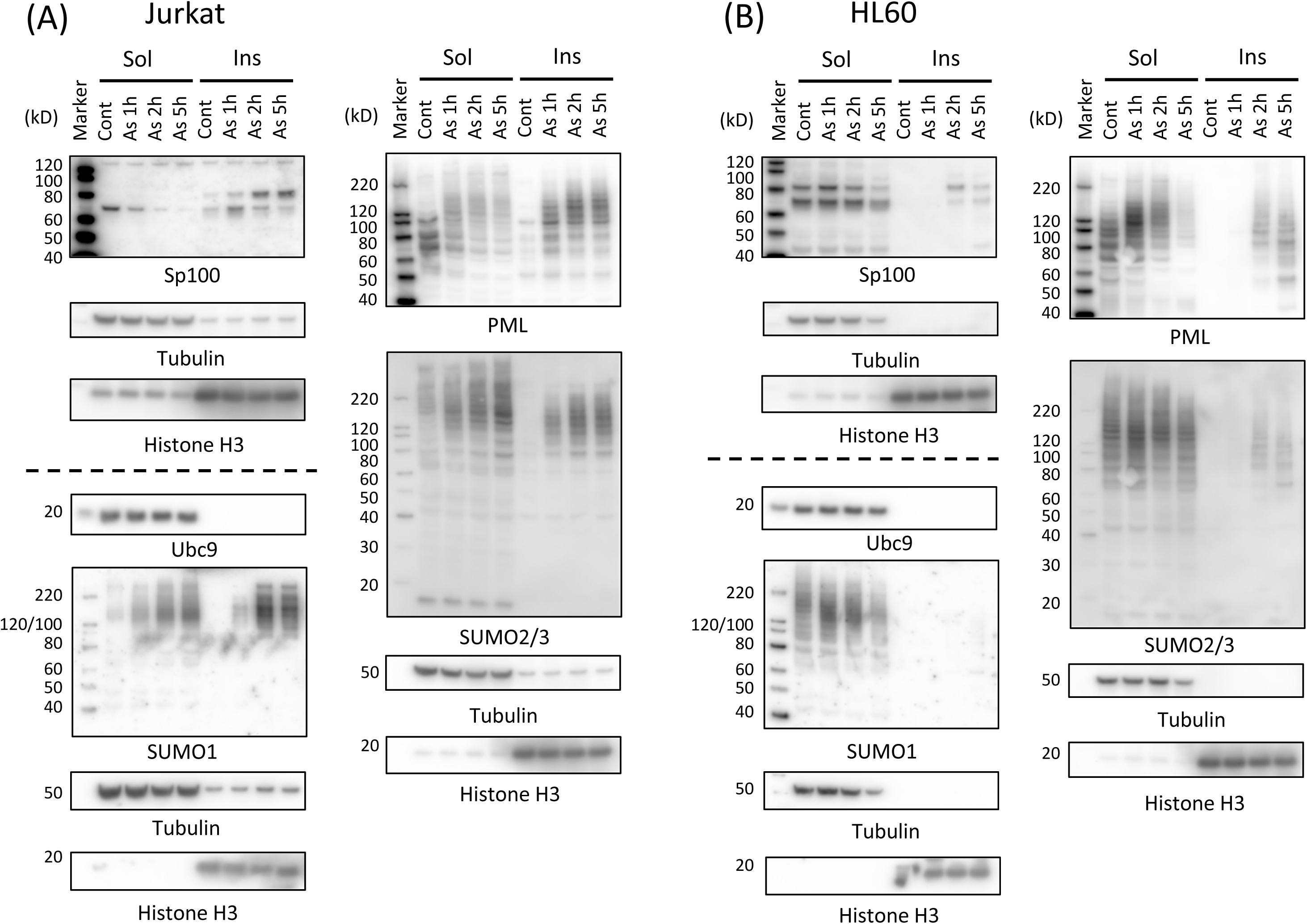
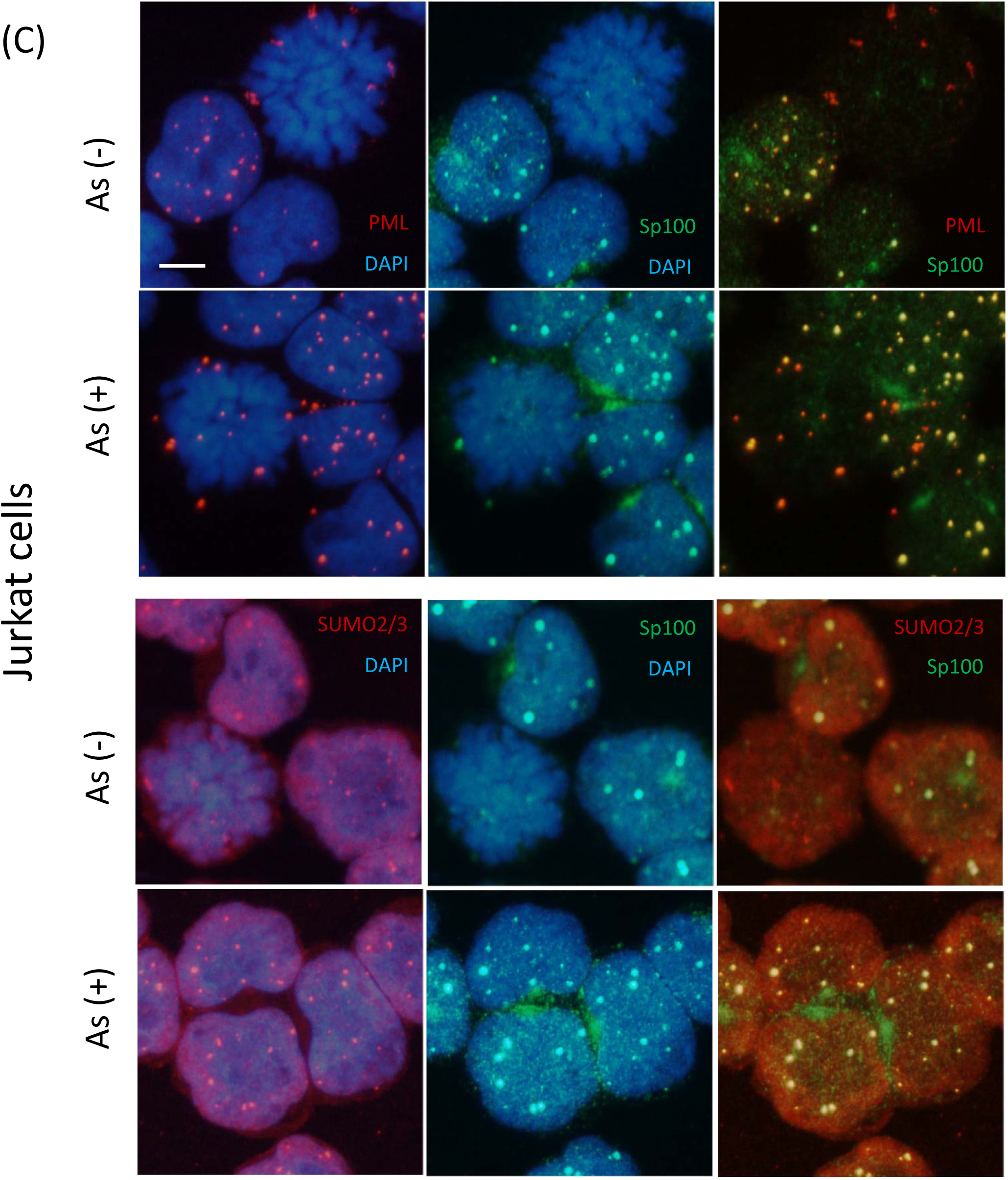
The solubility changes and SUMOylation of Sp100 and those of PML in Jurkat (A) and HL60 cells (B), and immunostaining of Jurkat cells with anti-SP100, anti-PML, and anti-SUMO2/3 antibodies (C). (A and B) The cells were cultured in suspension in 10% FBS-containing RPML1640 medium and exposed to 3 μM As^3+^ for 0 (Cont), 1, 2, and 5 h. The cells were centrifuged and lyzed with RIPA buffer. The soluble (Sol) and insoluble (Ins) fractions were analyzed for the detection of Sp100, PML, SUMO1, SUMO2/3, and Ubc9 by western blotting. (C) Jurkat cells were exposed to 3 μM As^3+^ for 2 h or left untreated. The cells were cytocentrifuged, fixed, and immunostained with anti-Sp100, anti-PML, and anti-SUMO2/3 antibodies. The nuclei were counterstained with DAPI. Scale bar = 10 µm.

**Fig. 9.**
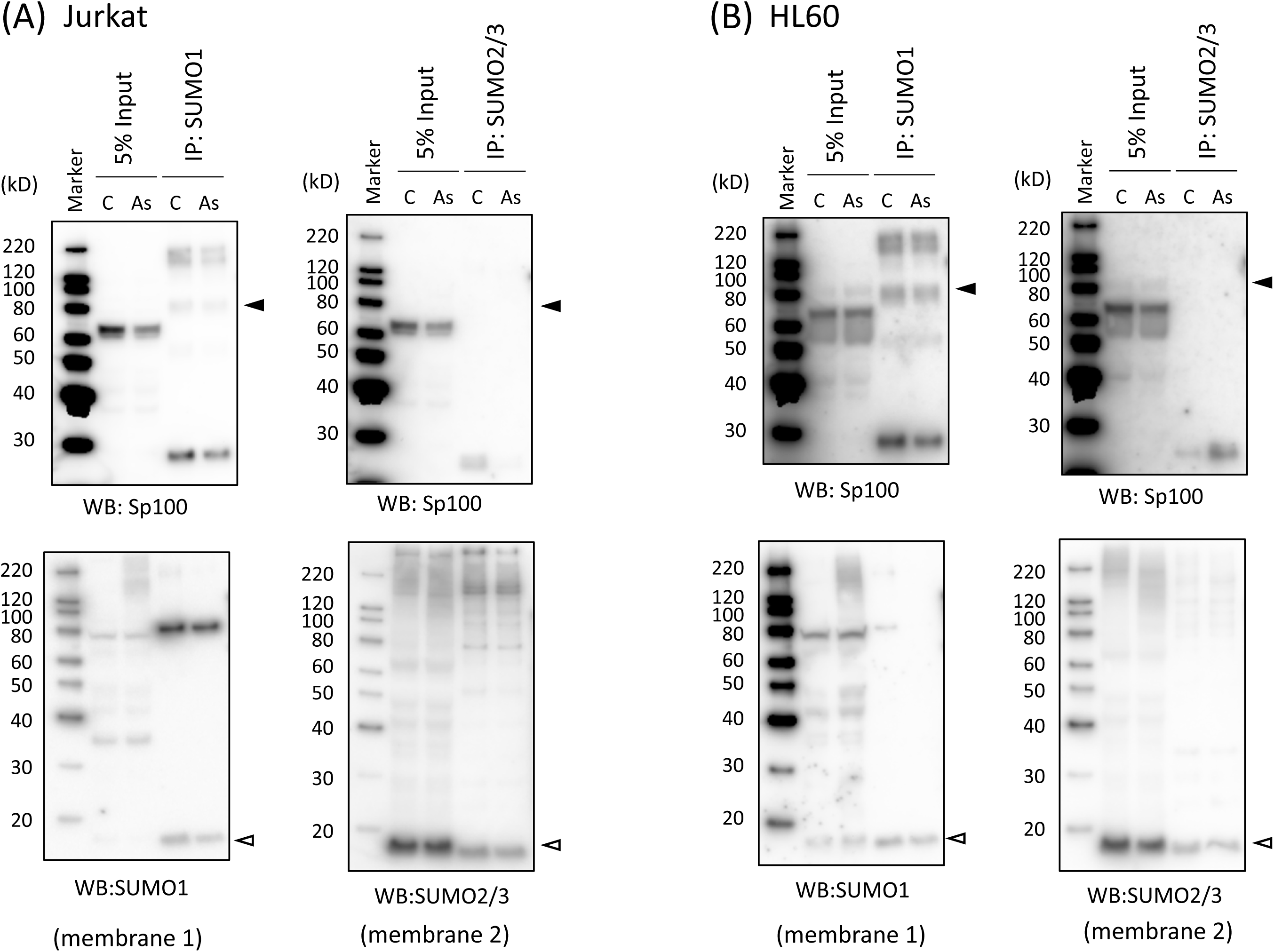
Immunoprecipitation with anti-SUMO1 and anti-SUMO2/3 antibodies to characterize SUMOylated Sp100 in Jurkat (A) and HL60 cells (B). The cells were treated with 3 μM As^3+^ (As) or left untreated (C) for 1 h and lyzed with RIPA buffer. The clear supernatant of the lysate was used for immunoprecipitation with SUMO1 and SUMO2/3 affinity beads. Closed triangle, SUMOylated Sp100. Open triangle, SUMO monomers.

## Discussion

### Responses of PML and SUMO to As^3+^

Endogenous PML isoforms in HEK293 cells and the overexpressed PML-VI in HEKPML cells responded to As^3+^ similarly. Both PML and SUMOylated PML were shifted from the RIPA-soluble to the RIPA-insoluble form by As^3+^, which is consistent with the notion that As^3+^ reinforces the PML-NB scaffold by oligomerization of PML [41]. Immunoprecipitation assay was performed using HEK293 and HEKPML cells stably expressing mCherry-tagged SUMO2 to study As^3+^-induced association of PML with SUMO. Unexpectedly, endogenous PML and SUMOylated PML were immunoprecipitated in mCherry-tagged ΔGG SUMO2-expressing HEK293 cells after exposure to As^3+^ at the comparable level in mCherry-tagged wild SUMO2-expressing HEK293 cells (Fig. 4B). These results indicate that As^3+^ induced non-covalent binding of PML proteins to SUMO2. However, DDK (FLAG)-tagged PML-VI was not pulled down by anti-RFP antibody in ΔGG mCherry-tagged SUMO2-expressing HEKPML cells even after exposure to As^3+^ (Fig. 5B), indicating that PML-VI did not associate with SUMO2 non-covalently. Together, our results show that As^3+^ strengthens SUMO2-PML association via the SUMO-SIM interaction because PML-VI is the only nuclear PML which lacks the SIM motif [42]. However, the SUMO-SIM interaction is required neither for the As^3+^-induced solubility change nor the covalent SUMOylation of PML because PML-VI responds to As^3+^ similarly to the other endogenous PML [32–34, 37]. To our best knowledge, this is the first report showing that As^3+^ induces non-covalent binding of SUMO to PML via the SUMO-SIM interaction.

It should be emphasized that SUMO2/3 molecules co-localized with PML in PML-NBs in both untreated and As^3+^-exposed cells (Fig. 1B), implying that the microscopically observed co-localization of PML with SUMO2/3 does not necessarily reflect covalent SUMOylation of PML [37]. However, inhibition of SUMO E1 enzyme with ML792 prevented SUMO2/3 from residing in PML-NBs irrespective of exposure to As^3+^ (Fig. 2), suggesting that the SUMO activation process is requisite for not only SUMOylation of PML, but also localization of SUMO in PML-NBs. These results may indicate that As^3+^ alters the SUMOylation-deSUMOylation balance on PML in favor of SUMOylation in PML-NBs. In addition, As^3+^ may activate the SUMO E3 ligase activity of PML, since it has been shown that PML increases the basal SUMOylation levels of KAP1 and DPPA2 and As^3+^ enhances SUMOylation of these proteins with SUMO2 in murine embryonic stem cells [43].

### Responses of DAXX and Sp100 to As^3+^

It is known that both DAXX and Sp100 are PML-NB client proteins and co-localize with PML in the nuclei [33, 44]. The present study also shows that Sp100 co-localized with PML and SUMO2/3 in Jurkat cells (Fig. 8). Thus, it is reasonable to suppose that these proteins may affect the responses of PML to As^3+^ and vice versa.

The involvement of PML in expression levels of DAXX is controversial. The protein level of DAXX is similar between wild-type and *Pml*^-/-^ MEF cells [23]. The depletion of PML diminishes both protein and mRNA levels of DAXX in HeLa cells, which suggests that DAXX is stabilized by PML [45]. In the present study, however, DAXX was significantly suppressed in HEKPML cells compared to HEK293 and *PML^-/-^* cells. These results warrant a further study to elucidate how DAXX is recruited to PML-NBs and stabilized by PML isoforms. The As^3+^-induced solubility shift and SUMOylation, which are hallmark responses of PML to As^3+^, were not observed in DAXX (Fig. 7 and Supplementary Fig. 3).

Heterochromatin protein 1 α (HP1α), a DNA-binding protein, resides at PML-NBs, but only via Sp100 [46]. It has been reported that As^3+^ induces SUMOylation of Sp100, which is delayed by PML silencing in H1299 and HeLa cells. [26]. The knockdown of PML has also been reported to decrease the SUMOylated form of Sp100 in THP-1 cells [44] and also in human fibroblasts [47]. The present study shows that expression and SUMOylation of Sp100 depend on the cell types. Sp100 was not detectable by our western blot method in HEK cells (Supplementary Fig. 2A). SUMOylation of Sp100 was enhanced by As^3+^ with the clear solubility shift in Jurkat cells, while a substantial amount of Sp100 was SUMOylated in the soluble fraction of untreated HL60 cells (Fig. 8). It is of interest to note that As^3+^-induced solubility shifts of both Sp100 and PML were less robust in HL60 cells than in Jurkat cells (Fig. 8).

The tripartite (RBCC) motif is critical not only for the formation of PML-NBs, but also for the As^3+^-induced solubility change and SUMOylation of PML [33]. 3KR mutated PML, in which SUMOylation is incompetent, fails to recruit Sp100, DAXX, and SUMO1 into PML-NBs [48]. DAXX and Sp100 do not co-localize and instead form discrete foci in PML-depleted human fibroblasts, while both DAXX and Sp100 converge on PML-NBs in wild-type cells [47]. These results indicate that DAXX and Sp100 reside in PML-NBs via PML-associated SUMO. Both DAXX and Sp100 have a least one SIM motif and neither DAXX nor Sp100 has the tripartite motif [25–27]. Our present results showed that As^3+^ increased the non-covalent association of PML with SUMO via the SUMO-SIM interaction besides the covalent SUMOylation of PML (Figs. 4 and 5), indicating that PML plays a pivotal role in accommodation of SIM-containing client proteins via SUMO in PML-NBs.

## Conclusions

In summary, we have provided an insight into As^3+^-induced changes of PML-NB client proteins. Sp100 and PML responded similarly to As^3+^ regarding the solubility and SUMOylation. Our current findings about the cell-type dependent expression of Sp100 and As^3+^-induced monoSUMOylation of Sp100 warrant a further study for the role of Sp100 in PML-NBs. The present study also highlights As^3+^-induced non-covalent association of PML with SUMO via the SUMO-SIM interaction. However, the SUMO-SIM interaction is not required for the As^3+^-induced solubility change or “stiffness” of PML-NB scaffolds and covalent SUMOylation of PML.

## Supporting information

This article contains supporting information. (Supplementary figures, and a source table for reagents)

## Acknowledgment

The authors would like to thank Ms. Mihoko Tadano for her technical assistance.

## Author contributions

SH designed the study, performed the experiments, and drafted the paper. OU and AKU performed the experiments and revised the paper.

