## Supplementary figures for "Arsenic induces two different interaction modes of SUMO with promyelocytic leukemia (PML) proteins"

**Supplementary figure legends**

**Supplementary Fig. 1, mRNA levels of *DAXX* in HEK293 (293), *PML*$^{-/-}$, and HEKPML cells.** HEK293, *PML*$^{-/-}$, and HEKPML cells were cultured to sub-confluence in quadruplicate. mRNA levels of *DAXX* and *GAPDH* were measured by RT-qPCR using the ΔΔCt method. The primer information is available from Supporting information. *, Significantly different from the other two groups (N=4).

**Supplementary Fig. 2, Expression levels of Sp100 (A) and RanGAP1 (B) in HEK293 (293), *PML*$^{-/-}$, and HEKPML cells.** See also the legend to Supplementary Fig. 1. (A) Neither Sp100 nor modified Sp100 was detectable in HEK cells by the current western blot analysis method. (B) Modification of RanGAP1 with SUMO1 occurred normally in HEK293 (293), *PML*$^{-/-}$, and HEKPML cells. The membrane was re-probed with anti-RanGAP1 (rabbit polyclonal) after probing with anti-DAXX (mouse monoclonal), and before the second re-probing with HRP-tagged anti-α-tubulin and HRP-tagged anti-histone H3. Thus, see Fig. 7 for the second re-probing with HRP-conjugated anti-tubulin and anti-histone H3 antibodies.

**Supplementary Fig. 3, Upregulation of Sp100 and PML by IFN in HL60 cells.** HL60 cells were cultured for 0, 1, and 3 days with 100 U/mL IFNα2a, and then exposed to 3 μM $As^{3+}$ or left untreated for 2 h. The soluble (Sol) and insoluble (Ins) fractions of the cell lysates were used for the detection of Sp100, PML, and DAXX. The membrane was first stained with Ponseau S solution.

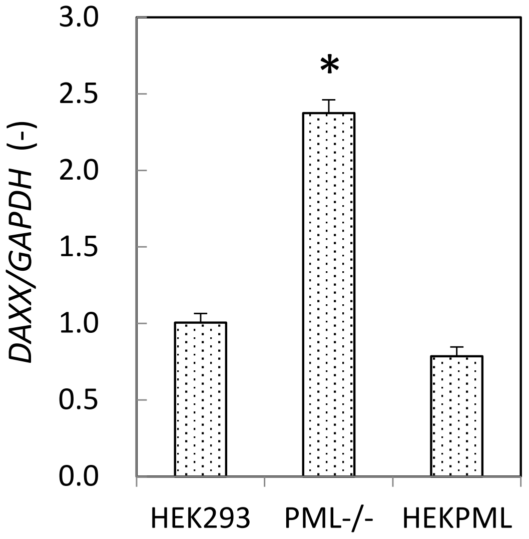


Supplementary Fig. 1

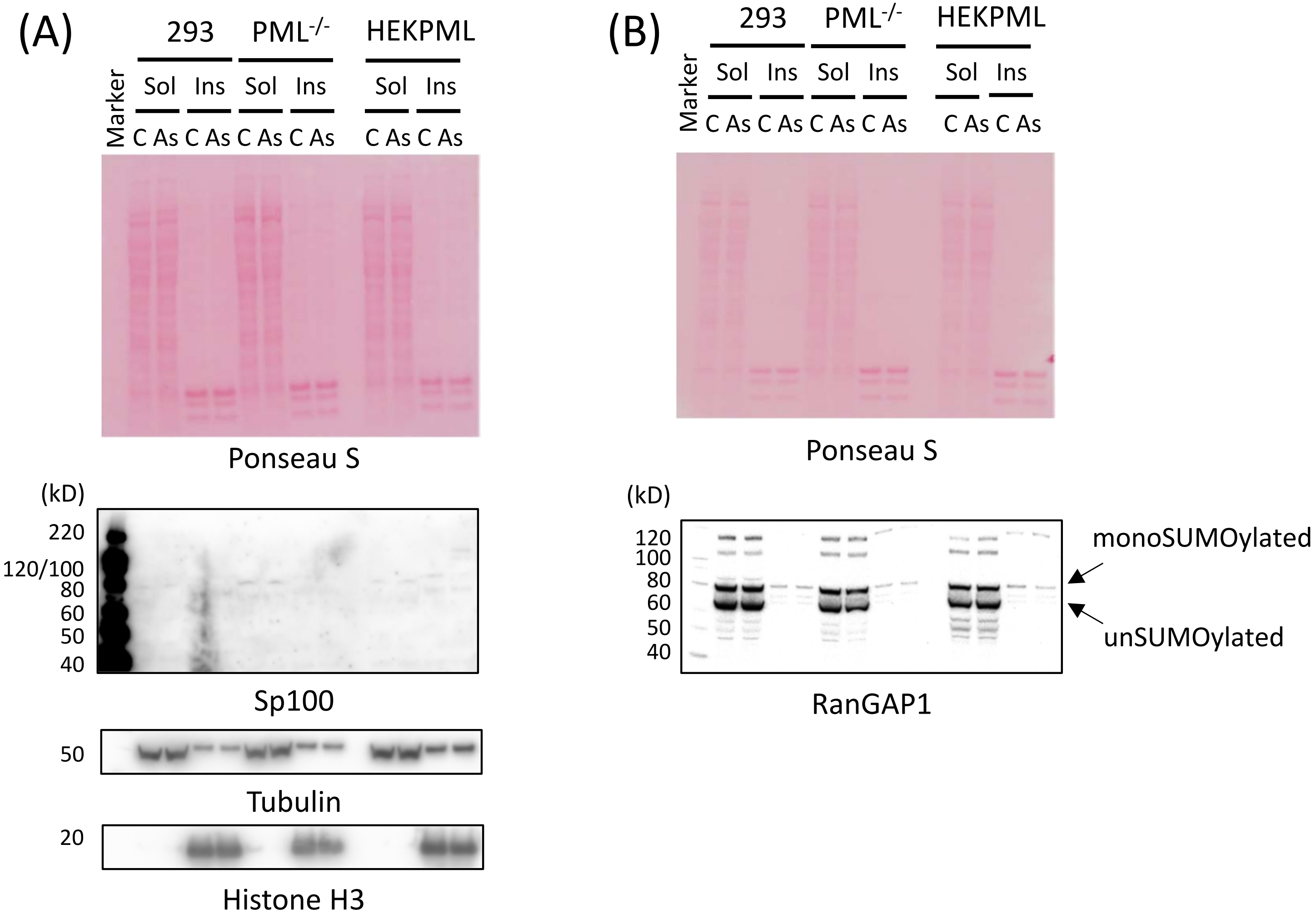


Supplementary Fig. 2

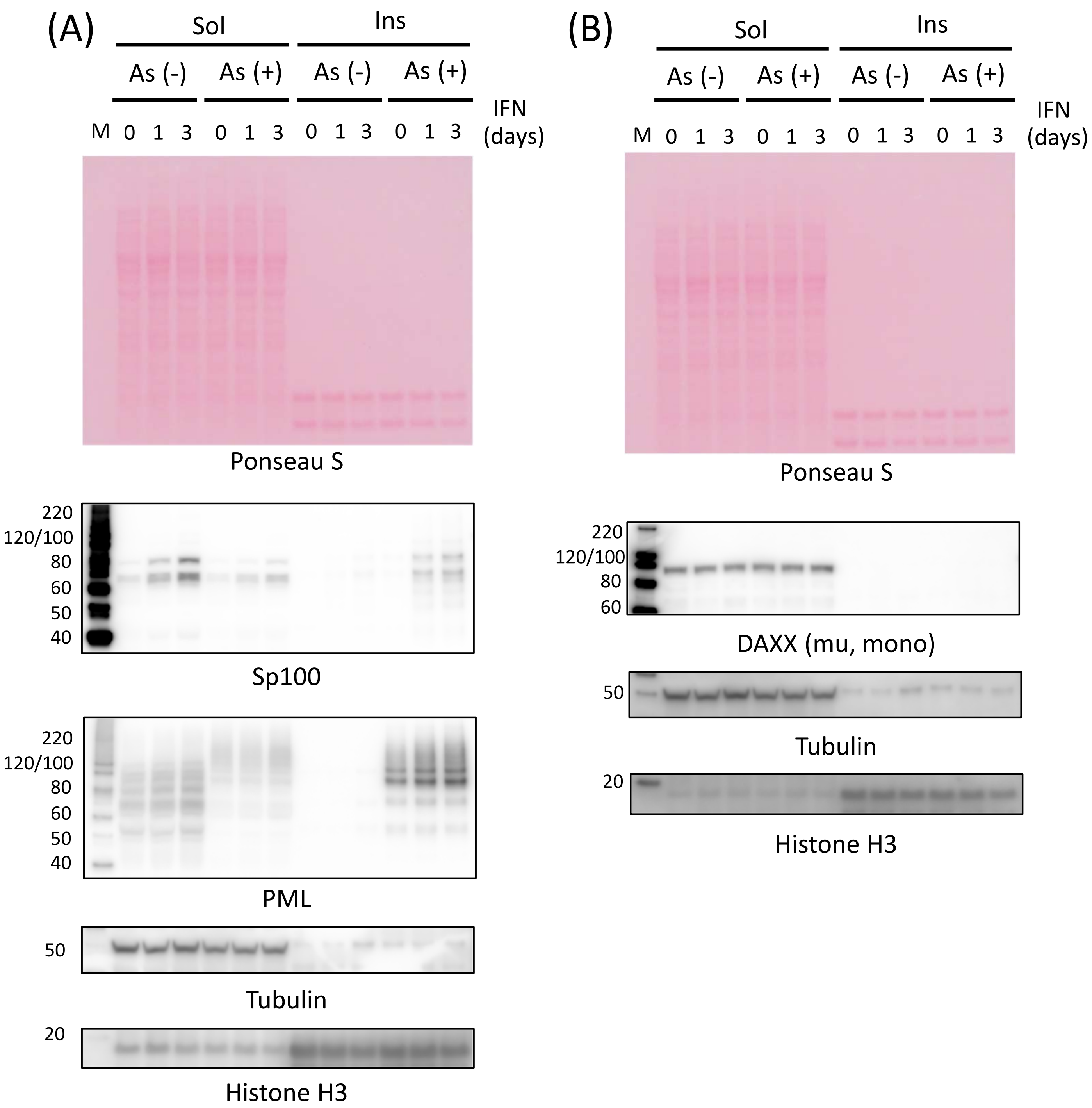


Supplementary Fig. 3
